# Short-term thinning effect on inter- and intra-annual radial increment of Mediterranean Scots pine-oak mixed forest

**DOI:** 10.1101/2023.06.22.546107

**Authors:** J. Aldea, M. del Río, N. Cattaneo, J. Riofrío, C. Ordóñez, S. Uzquiano, F. Bravo

## Abstract

Thinning treatment and mixed forest stands have been suggested as possible adaptation strategies to cope to climate change but there is still scarce knowledge about the combination of both subjects. In this study, we aim to better understand the thinning effect and the growth differences between two coexisting species on inter- and intra-annual cumulative radial increment patterns. We studied radial increment of a Scots pine-oak (*Pinus sylvestris-Quercus pyrenaica*) Mediterranean mixed forest during two climatically contrasted years (2016-2017) in north-western Spain. Data came from a thinning trial consisting in a randomized block experimental design with a control and two thinning treatments from below: a moderate and heavy thinning removing 25% and 50 % of initial basal area respectively focused on both species. Tree radial increment was analyzed based on bi-weekly readings from band dendrometers installed in 90 oak and pine trees. Non-linear mixed model based on double-Richards curve was fitted to show thinning and species differences in intra-annual cumulative radial increment patterns. Inter-annual basal area increment at species and stand levels were estimated using the model previously fitted at tree level and aggregating the results for exploring thinning effects at these levels. Scots pine leaded the tree and stand growth, and had also a better respond to early spring drought compared to oak. Heavy thinning increased tree radial increment for both species at the expense of decreased stand basal area. At species level, basal area increment decreased for Scots pine, however, heavy thinning generated the same oak basal area increment than control. Thus, heavy thinning may be good strategy towards a conversion from overaged coppice stands into high forests to conform a stable mixed forest stand.

**Highlights:**

- Scots pine leaded growth at tree and stand level
- Scots pine trees may take advantage during early spring droughts
- Heavy thinning increased tree radial increment for both species
- Heavy thinning decreases stand basal area growth

## 1. Introduction

Climate change represents one of the greatest threat to forest vitality and, consequently, a challenge for managers is to develop adaptive strategies that minimize ecosystem vulnerability (Allen et al., 2010). Several studies have already evidenced decreases in forest productivity and increased tree mortality or dieback in different regions of the world, especially in southern Europe (Calama et al., 2023; Camarero et al., 2018; Galiano et al., 2011; Martínez-Vilalta et al., 2012). Some strategies as changes in forest structure and/or species composition have been recommended to reduce the impacts of climate change (D’Amato et al., 2013).

Competition affects the way of the tree growth response to climate (Castagneri et al., 2022; Fernández-de-Uña et al., 2015; Sánchez-Salguero et al., 2015). Thus, forest management techniques such as thinning has been suggested to promote climatic resilience in forest stands by moderating tree competition and changing stand structure (Moreau et al., 2022; Olivar et al., 2014; Sohn et al., 2016a, 2016c). Residual trees gain increased access to environmental resources (i.e., soil moisture, light), promoting the vigor of individual trees and release stress from climatic conditions (Magruder et al., 2013). Tree response to thinning treatments depends on the thinning regime (age at the first thinning, intensity, type, and frequency). Heavy thinning, is a suitable approach to improve the growth response of remaining trees to drought in both conifers and broadleaves (Sohn et al., 2016c). However, this effect may decrease over time since the last thinning and could even become negative when compared to unthinned stands of *Pinus sylvestris* L. (Sohn et al., 2016a, 2013). Reduced competition through thinning was also found to stimulate and and prolong the duration of the growing season for beech (*Fagus sylvatica* L.) regardless weather conditions (van der Maaten, 2013). The reduction of stand density increases drought resistance and resilience in xeric Scots pine forests, thus, may decrease stand vulnerability to drought (Giuggiola et al., 2013). Similarly, thinning treatments have been shown to increase growth, reduce long-term stress caused by competition for water, and to reduce the vulnerability of trees to drought in Mediterranean environment (del Río et al., 2017b; Navarro-Cerrillo et al., 2019).

Mixed forest stands has been suggested as a possible adaptation strategy to counteract climate change, increasing resilience to biotic and abiotic factors (Grossiord, 2019; Guyot et al., 2016; Jactel et al., 2017; Pardos et al., 2021) and enhancing the level and temporal stability of productivity (Bauhus et al., 2017a; del Río et al., 2022, 2017b). They can also present some advantages compared to monospecific stands with regard to ecological functions and services (Felton et al., 2016; Huuskonen et al., 2021). Thereby, combining mixed forest and thinning effects as feasible adaptation strategies to cope with climate change could contribute to promote social, economic and environmental functions towards an adaptive forest management. Nevertheless, there is few information about thinning effects on the resilience of mixed forest stands (Navarro-Cerrillo et al., 2016; Vernon et al., 2018).

Radial increment analysis is commonly used to estimate tree growth and to evaluate the ecological consequences and production alterations motivated by climate change or management interventions (Siegmund et al., 2016). Understanding stem growth differences of co-existing species led to assess stand stability on aboveground wood production and predict change in species composition (Jucker et al., 2014). Intra-annual radial increment investigation permits evaluate tree growth species differences in response to particular climatic events that may escape a classical inter-annual dendroclimatic approach (Duchesne and Houle, 2011). However, intra-annual radial variation studies focused on mixed forests in the Mediterranean region are still scarce (Aldea et al., 2018; de-Dios-García et al., 2018).

Here, we evaluated species differences in intra- and inter-annual radial increment patterns of *Pinus sylvestris* L. and *Quercus pyrenaica* Willd. (hereinafter Scots pine and oak, respectively) in thinned and unthinned mixed stands in Mediterranean climate. Both species form spontaneous mixed stands where their natural distribution areas coincide. Consequently, forest management strategies during the second half of the twentieth century included re-introducing pine species into oak coppice stands as a method to increase soil protection and stand productivity. Scots pine is a typical pioneer and light-demanding species with a fast radial growth (Sánchez-Costa et al., 2015). On the other hand, deciduous and more shade-tolerant oak trees would utilize resources more efficiently, so it predominates in the late-successional stage (Cuny et al., 2012; Rodríguez-Calcerrada et al., 2010). These differences in functional traits and growth strategies could decrease competition between species, and increase the supply, capture, or use efficiency of site resources (Forrester, 2017, 2014; Grossiord, 2019). Therefore, Scots pine-oak mixed stands can support higher volume increment per occupied area compared to pure stands (del Río and Sterba, 2009; Steckel et al., 2019). This spatial and temporal complementarity could also cause, consequently, a climate sensitivity reduction (Merlin et al., 2015; Steckel et al., 2020; Toïgo et al., 2015) and a more stable productivity (del Río et al., 2022).

Although a large number of thinning experiments have been established for Scots pine in the last century across Europe (del Río et al., 2017a), thinning experiments in mixed stands are currently scarce (Bauhus et al., 2017c; Primicia et al., 2016). Heavy thinning is commonly associated to a yield loss at stand level for pure Scots pine stands (del Río et al., 2017a). In contrast, they may increase growth at tree and stand levels for oak coppice stands (Cañellas et al., 2004; Corcuera et al., 2006). Cotillas et al., (2009) also found that selective thinning (removing 20–30% of total stump basal area) improved tree growth for oak mixed coppice stands under natural and reduced rainfall conditions. Aldea et al., (2017), analyzing intra- and inter-annual radial increment, revealed a positive thinning effects on growth (regardless drought conditions) in Maritime pine and oak mixed stands. However, growth response to thinning in Scots pine-oak mixed forests in Mediterranean area has not been yet studied. In spite of the wide forest cover area and ecological and socio-economic importance of this mixture, there is not a specific silviculture for mixed stands of this species composition.

We used band dendrometer measurements over two climate contrasted years (2016 and 2017) from a thinning trial to show species differences and short-term thinning effects on inter- and intra-annual cumulative radial increment patterns. The aims of the study were: 1) to decipher species growth differences; and 2) to quantify the short-term thinning effect at different spatio-temporal scales (tree vs. species and stand levels, and intra-vs. inter-annual levels). We tested the hypotheses that (i) pine leads tree and stand growth and that (ii) heavy thinning increases radial increment at tree level, but implies a reduction of stand production for both species at short time.

## 2. Materials and methods

### 2.1. Study site

The experiment was located in Palacio de Valdellorma (León, 42°45’42.4’’N, 05°12’39.6’’W) in north-western Spain. The experimental site was sited at 990 m.a.s.l. in a continental Mediterranean climate. The average annual rainfall is 515 mm with a marked summer drought period between July and August, when 42 mm of precipitation are usually recorded (AEMET, 2018, Spanish State Meteorological Agency. 2661 weather station code, based on 1981-2010 historical records). Annual mean temperature is 11.1 °C and the hottest month is July, with an average temperature of 27.4 °C. The probability of frost period (temperature below 0 °C) is from December to February. Topography was moderate with a slope of 16% and soils consisted in acid conglomerates based on Miocene clay sediments (IGN, 1991).

Initially, an original oak coppice stand was harvested during 1970’s and a reforestation was carried out by planted Scots pine in rows. Oak coppice sprouts grew again via asexual reproduction between pine rows, so at experimental establishment date the stand looked as pine-oak even-aged mixed stand of 40 years old, although real cambial age differed between species. Stand basal area was dominated by pine, with a mean species proportion in basal area of 70%.

Nine rectangular plots (50 x 40 m) were established following a complete randomized block design. The experimental design consisted of two thinning treatments with different intensity and unthinned control (C) with three replicates each. Thinning treatment consisted in moderate (MT) and heavy thinning (HT) removing 25% and 50 % of initial basal area, respectively (Table 1). Trees of both species were removed applying thinning from below, which involved logging the suppressed and intermediate trees. Species proportion by plot was approximately maintained, which results in 60-70 % of Scots pine and 30-40% of oak in the remain basal area. Thinning was carried out at winter of 2015 and the felled logs and branches were removed from plots. In each plot, the diameter at breast height (DBH) of all trees were measured whereas tree height was measured in a subsample covering the diameter distribution by species. Mean DBH, stand density, dominant height and stand basal area were estimated by species. There were no statistical differences between plots for each species before thinning. As expected, after thinning statistical differences were identified in mean DBH, stand density and basal area according to Tukey contrast (Table 1).

**Table 1.** Main stand characteristics before and after thinning for each species. Data shown are mean and standard deviation values by treatment. Different letters denote significant differences after thinning at the 0.05 significance level. H_0_: dominant height; dbh: mean diameter at breast height; BA: stand basal area; C: unthinning; MT: moderate thinning (25% BA removed); HT: heavy thinning (50% BA removed)

| Species | Treatment | $H_0$ (m) | Before thinning | | | After thinning | | |
| --- | --- | --- | --- | --- | --- | --- | --- | --- |
|  |  |  | Density<br>(n·ha<sup>-1</sup>) | dbh<br>(cm) | BA<br>(m<sup>2</sup>·ha<sup>-1</sup>) | Density<br>(n·ha<sup>-1</sup>) | dbh<br>(cm) | BA<br>(m<sup>2</sup>·ha<sup>-1</sup>) |
| **Pinus**<br>**sylvestris** | **C** | 12.5±1.2 | 1,415±130 | 12.2±4.3 | 19.6±2.3 | 1,415±130c | 12.2±4.3a | 19.6±2.3b |
|  | **MT** | 10.7±1.9 | 1,575±90 | 12.0±4.5 | 19.5±3.9 | 710±240b | 16.0±3.3b | 13.7±5.5b |
|  | **HT** | 11.7±1.7 | 1,580±130 | 11.8±4.2 | 20.9±3.3 | 390±197a | 15.1±3.5b | 9.0±4.6a |
| **Quercus**<br>**pyrenaica** | **C** | 10.9±1.1 | 2,960±710 | 5.9±3.2 | 12.6±1.6 | 2,960±710b | 5.9±3.2a | 12.6±1.6b |
|  | **MT** | 10.7±0.7 | 2,855±366 | 6.6±3.5 | 12.1±1.4 | 1,195±286a | 9.2±4.6b | 8.8±1.7a |
|  | **HT** | 11.3±1.8 | 2,005±760 | 6.6±4.1 | 8.7±5.0 | 430±427a | 9.3±6.3b | 4.0±2.9a |

### 2.2. Stem radial variation and climatic measurements

We installed bands dendrometer (DB 20, EMS Brno) after thinning treatments on five trees per species and plot, according to a stratified sampling approach that took diameter distribution into account. A total of 90 dendrometer bands were fitted at breast height (1.30 m) after partial bark removal (rhytidome) to reduce stem rehydration effect. Band dendrometers were measured every two weeks through-out the year to the nearest 0.1 mm from April 2016 to December 2017. All measurements were taken in the morning to reduce diurnal bias, which is caused by stem shrinkage from transpiration. The measurements were corrected for temperature effects and dendrometer thermal expansion following the manufacturer’s instructions (11.2 × 10^−6^mm◦C^−1^). Finally, girth increment data were transformed to radial increments based on a hypothetic cylindrical tree shape.

The weather during sampling years (2016-2017) was characterized by summer drought (Figure 1). However, 2017 had a severe drought due to a low precipitation at the beginning of spring (March-April) and during autumn, prolonging the drought period until the end of autumn (October). Accordingly, annual precipitation was 515 mm and 368 mm for 2016 and 2017 respectively. The average of daily maximum temperatures for warmest month was higher than mean historical records (29.6 °C and 28.6 °C for 2016 and 2017 respectively vs. 27.3°C for 1981-2010 period). Monthly temperature and precipitation records were compiled using data from the closets AEMET automatic network stations (León, Sahechores, 2626Y, sited at 25 km from the study site).

**Figure 1.**
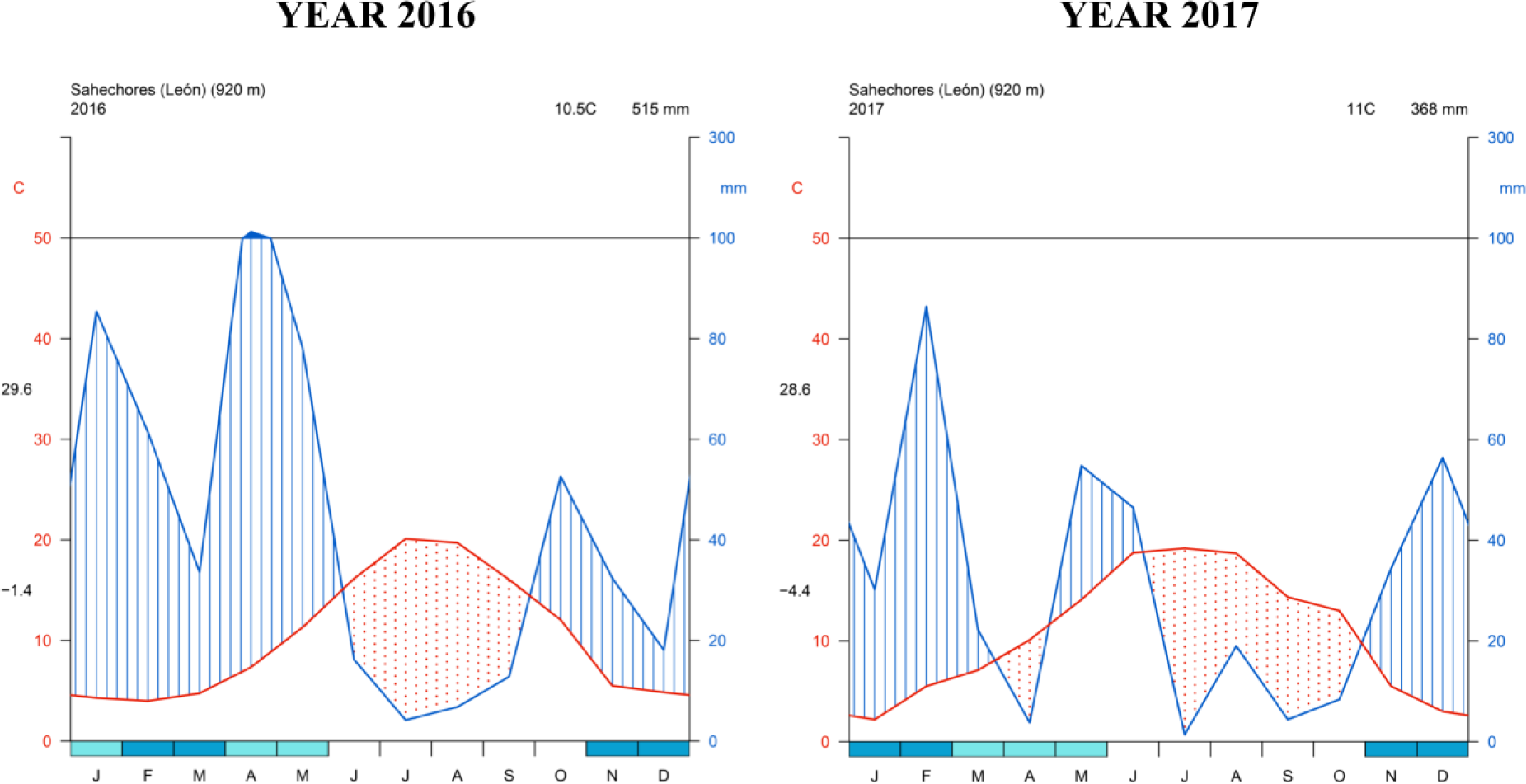
Walter & Lieth climograms for sampled years. Digits sited on the left side of y-axis are average of daily maximum temperatures of warmest month and average of daily minimum temperatures of coldest month from top to bottom respectively. Source: AEMET, (2018).

### 2.3. Data analysis

#### 2.3.1. Intra-annual analysis at tree level

Cumulative radial increment patterns of both species showed clearly bimodal pattern, which is typical of Mediterranean environments (Albuixech et al., 2012; Aldea et al., 2017; Pacheco et al., 2018): stem growth in spring season, contraction during the summer due to depletion of stored water (concurrent with increasing water deficit) and stem rehydration (with growth for certain species) after autumn rainfall. Because of that, we employed a non-linear mixed model based on the sum of two Richard curves (double-Richards) to consider the bi-modal pattern of intra-annual radial increment. The advantage of Richards curve compared to commonly used Logistic or Gompertz function is that a fourth parameter included in the former, allows a better and flexible fit to the original data, avoiding convergence problems. In fact, Logistic or Gompertz function could be considered as a concrete case of a generalist Richards curve (Oswald, 2015).

We determined the most appropriate model (i.e., number of necessary Richards curve parameters, see Eq.1) for our data comparing all possible positive-negative Richards non-linear models by ranking and comparing nested models using extra sum of squares F-Tests (Oswald, 2015).

A random effects structure was also included into the double-Richards model to consider the spatial and temporal dependence of measurements (measurement nested in tree, tree nested in plot). This structure was evaluated fitting a beyond optimal model with different random structures maximizing the restricted log-likelihood (Zuur et al., 2009) and then comparing and selecting the lowest value of the Akaike information criterion (AIC). Then, species, thinning treatment and year fixed effects (as categorical) and diameter covariate (continuous) were included to evaluate their effect on Richards curve parameters. We compared all possible fixed model structures including permutation in the input of the variables in the model, combination of variables affecting several parameters and interaction among them (e.g. different thinning effects by species or year). The final model was selected according the most parsimonious (low AIC) and biological sense.

Finally, serial autocorrelation was assessed by partial and autocorrelation function plots, and several serial correlation structures were evaluated (autoregressive, moving average, and a mixed autoregressive-moving average model) to model residual autocorrelation. A variance function for modeling heteroscedasticity was also used according several structures (exponential, power, and constant plus power of the absolute value of the variance covariate) (Pinheiro and Bates, 2000). Model fitting was performed with the ‘FlexParamCurve’ (Oswald, 2015) and ‘nlme’ (Pinheiro et al., 2015) packages in R (R Development Core Team, 2020).

#### 2.3.2. Inter-annual analysis at species stand level

The species stand basal area growth was estimated based on measured tree diameters distribution by plot and using the model fitted previously at tree level. It was carried out by the empirical best linear unbiased predictors (EBLUPs) for a given level of random variability (i.e. plot) predicted in a previous step along with different explanatory covariates acting at that level (e.g. species, thinning treatment, year and diameter) for all remaining trees after thinning. The basal area value per tree, assuming a cylindrical tree shape, was aggregated for each plot obtaining thus the stand basal area increment. This estimated basal area was up-scaled to hectare. Then thinning treatment differences by species and year were tested using “Tukey” contrasts based on global rankings, simultaneous confidence intervals and adjusted p-values. The R packages ‘sae’ (Molina and Marhuenda, 2015) and ‘nparcomp’ (Konietschke et al., 2015), were used for calculate EBLUPs and Tukey test respectively.

## 3. Results

### 3.1. Tree radial increment model with best structure

The best model structure resulted in a five parameter model (all parameters from first Richards curve and asymptote of second Richards curve) and three invariable parameters (inflection point, rate and shape parameter from second Richards curve). These invariable parameters took the mean value across all trees in the dataset, reducing complexity and computation, due to these parameters did not vary across group levels (trees from plots). Plot and tree (nested in plot) random effects were added to the model, affecting only the intercept of spring asymptote (A_ij_), which turned out to be the best random structure, according to the model:

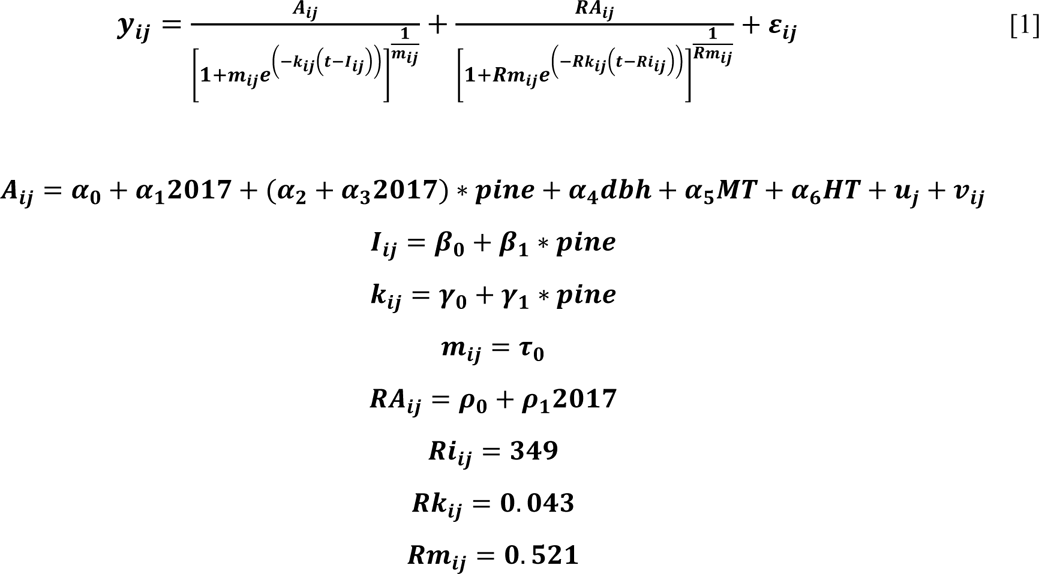

where ***y_ij_*** is the cumulative radial increment for tree i in plot j; ***α***_***i***_, ***β***_***i***_, ***γ***_***i***_ and ***τ***_***i***_are the asymptote, inflection point, rate and shape parameter regression of fixed effect variables for the first Richards curve and ***ρ***_***i***_ are the asymptote parameters for the second one (for clarification of parameters, see Figure S1); *2017* is a year dummy variable with value 0 for year 2016 and 1 for 2017; *pine* is also a dummy variable with value 0 for oak and 1 for pine species; *dbh* is tree diameter at breast height (mm) recorded before the thinning; *MT* is a dummy variable with value 1 for moderate thinning and 0 otherwise; *HT* is a dummy variable with value 1 for heavy thinning and 0 otherwise; ***u***_j_∼N(0, **σ**_***e***_) is the plot random effect; ***v***_ij_∼N(0, **σ**_ij_) is the tree random effect and ***ɛ***_ij_ ∼N(0, **σ**_***e***_) is the error term. ***Ri***_ij_, ***Rk***_ij_ and ***Rm***_ij_ are the inflection point, rate and shape invariable parameters from second Richards curve which took the mean value from double-Richards curve fitted for each tree (as mentioned previously).

Intra-annual cumulative radial increment patterns differed between species, thinning treatments and among years according to fitted double-Richards model (Table 2). Scots pine presented a higher increment rate (***γ***_**1**_), but earlier inflection point (***β***_**1**_) than oak regardless year. The inflection point was reached at 14 May and 16 June for Scots pine and oak species, respectively. Scots pine presented higher spring asymptotic value than oak for unthinned stands (***α***_**3**_ and Figure 2), although during 2016 asymptote difference between species was only caused by size differences (***α***_**2**_ and ***α***_**4**_). Year (climate) effect influenced both, spring and autumn asymptotes. Spring asymptote decreased in 2017 for oak trees (***α***_**1**_) due to spring drought but increased for Scots pine (***α***_**3**_) (Figure 2). Autumn asymptote increased in 2016 but with no statistical significance (***ρ***_**0**_). Both species were affected by autumn drought in 2017, so the autumn asymptotic value decreased significantly (***ρ***_**1**_). Exclusively heavy thinning increased significantly spring asymptotic value for both species (***α***_**6**_) (Table 2). However, moderate thinning could increase also tree radial increment for oak compared to unthinned stands (Figure 3), although it was not revealed in the fitted model.

**Figure 2.**
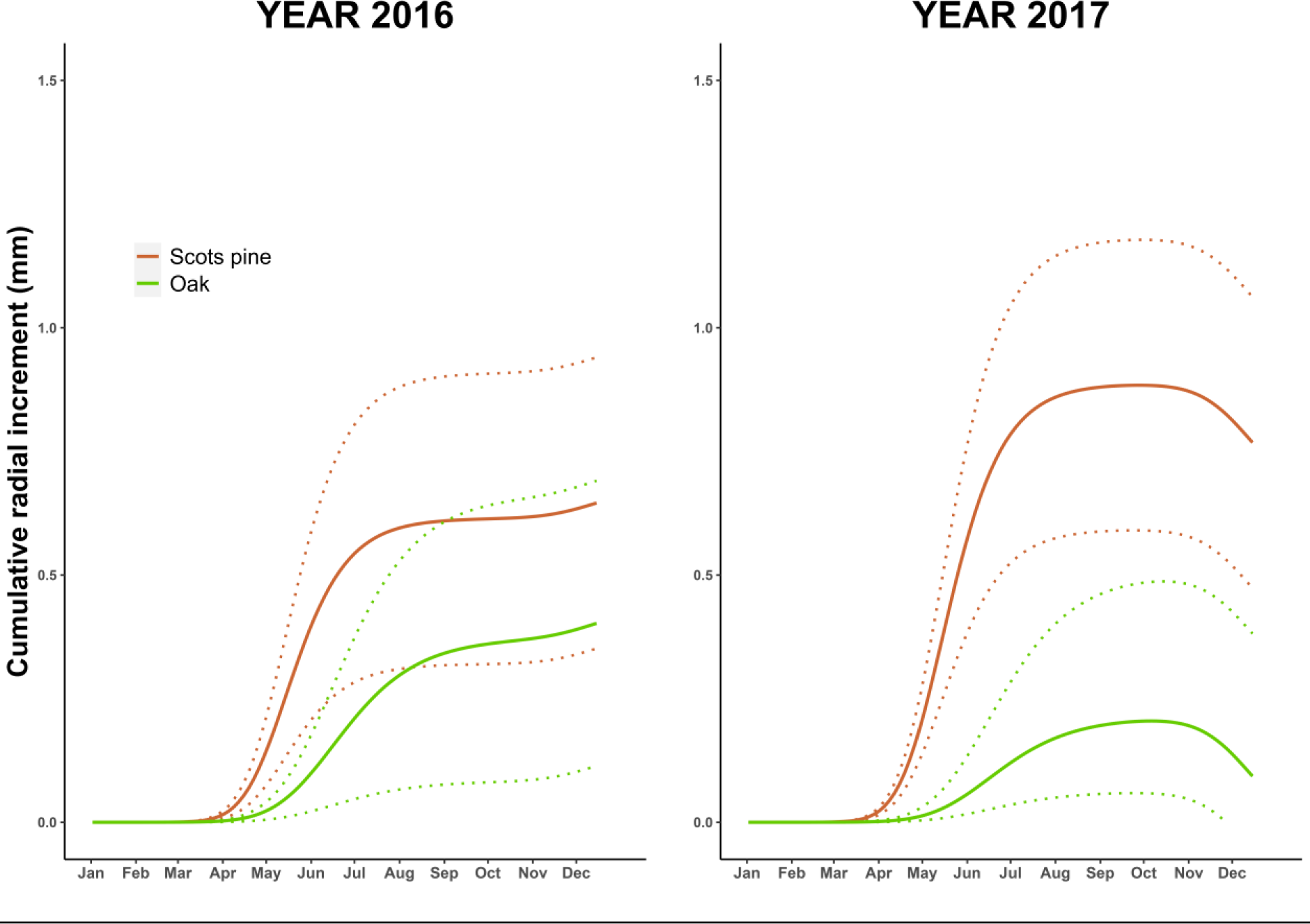
Species differences in intra-annual cumulative radial increment patterns calculated from the model predictions for the unthinned stands and both years. The thick solid lines represent double-Richards fitted models for Scots pine (red) and oak trees (green) for the species’ mean diameter. The dotted lines show values for a hypothetical tree with <u>±</u> 1 standard deviation from the species mean diameter.

**Figure 3.**
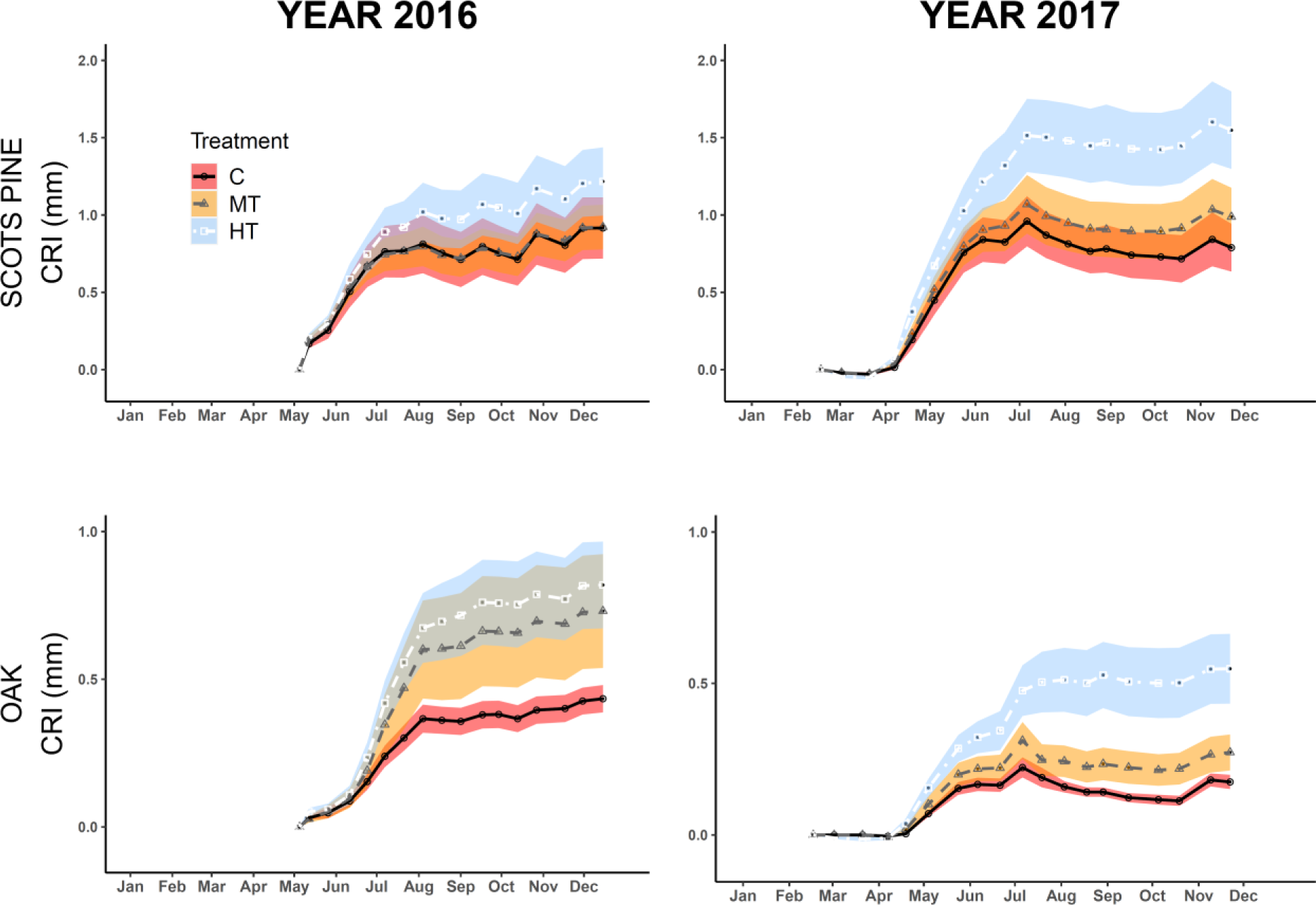
Unthinned and thinning treatment differences based on observed values of cumulative radial increment patterns of each year. Circles and solid line, triangles and dashed line, and square and dot-dashed line show mean pattern for trees in unthinned, moderate and heavy thinning stands, respectively. Shaded areas denote standard error: red for unthinned, orange for moderate and blue for heavy thinning remaining trees.

**Table 2.** Model fitted for intra-annual cumulative radial increment pattern (*Eq. 1*). Significant parameters are in bold. ***α***_***i***_, ***β***_***i***_, ***γ***_***i***_ and ***τ***_***i***_ are the asymptote, inflection point, rate and shape parameter regression of fixed effect variables for the first Richards curve and ***ρ***_***i***_ are the amplitude parameters for the second one; **σ**_j_ is standard deviation for plot random effect; **σ**_ij_ is standard deviation for tree random effect; **σ**_***e***_ is standard deviation for error term; **θ** is the residual serial correlation parameter for moving average model MA(1); **δ** is function parameter used to model residual variance as a power of the absolute value of the variance covariate (***g***_ij***k***_): **Var**(***ɛ***_ij***k***_) = **σ**^**2**^(|***g***_ij***k***_|^**δ**^)^**2**^

| Parameter | Coefficient | p-value |
| --- | --- | --- |
| $\alpha_0$ ($A_{2016}$) | -0.839 | <0.001 |
| $\alpha_1$ ($A_{2017}$) | -0.159 | <0.001 |
| $\alpha_2$ ($A_{2016\text{pine}}$) | 0.081 | 0.424 |
| $\alpha_3$ ($A_{2017\text{pine}}$) | 0.432 | <0.001 |
| $\alpha_4$ ($A_{\text{dbh}}$) | 0.101 | <0.001 |
| $\alpha_5$ ($A_{\text{MT}}$) | 0.124 | 0.303 |
| $\alpha_6$ ($A_{\text{HT}}$) | 0.374 | 0.002 |
| $\beta_0$ (I) | 167.274 | <0.001 |
| $\beta_1$ ($I_{\text{pine}}$) | -32.592 | <0.001 |
| $\gamma_0$ (k) | 0.032 | <0.001 |
| $\gamma_1$ ($k_{\text{pine}}$) | 0.012 | <0.001 |
| $\tau_0$ (m) | 0.307 | 0.032 |
| $\rho_0$ ($RA_{2016}$) | 0.070 | 0.065 |
| $\rho_1$ ($RA_{2017}$) | -0.337 | 0.002 |
| $\sigma_j$ ($A_{\text{plot}}$) | $2.16 \cdot 10^{-4}$ | |
| $\sigma_{ij}$ ($A_{\text{tree}}$) | 0.409 | |
| $\sigma_e$ (error) | 0.195 | |
| $\delta$ | 0.304 | |
| $\theta$ | 0.683 | |

### 3.2. Species stand basal area increment

The stand level analysis revealed that thinning treatments caused the lowest basal area increment for Scots pine regardless year (Figure 4). For oak, heavy thinning generated the highest basal area increment but without statistical differences with unthinned stands. Consequently, as Scots pine is the dominant tree species according to height and diameter stand structure, total stand basal area increment, i.e. for the sum of both species, decreased by thinning but without differences between the two thinning intensities. In addition, Scots pine stand basal area increment tended to be higher than oak, but with no differences for heavy thinning (P=0.101). Stand basal increment was stable along time, i.e., no significant (P>0.05) differences were found between years for each treatment and species.

**Figure 4.**
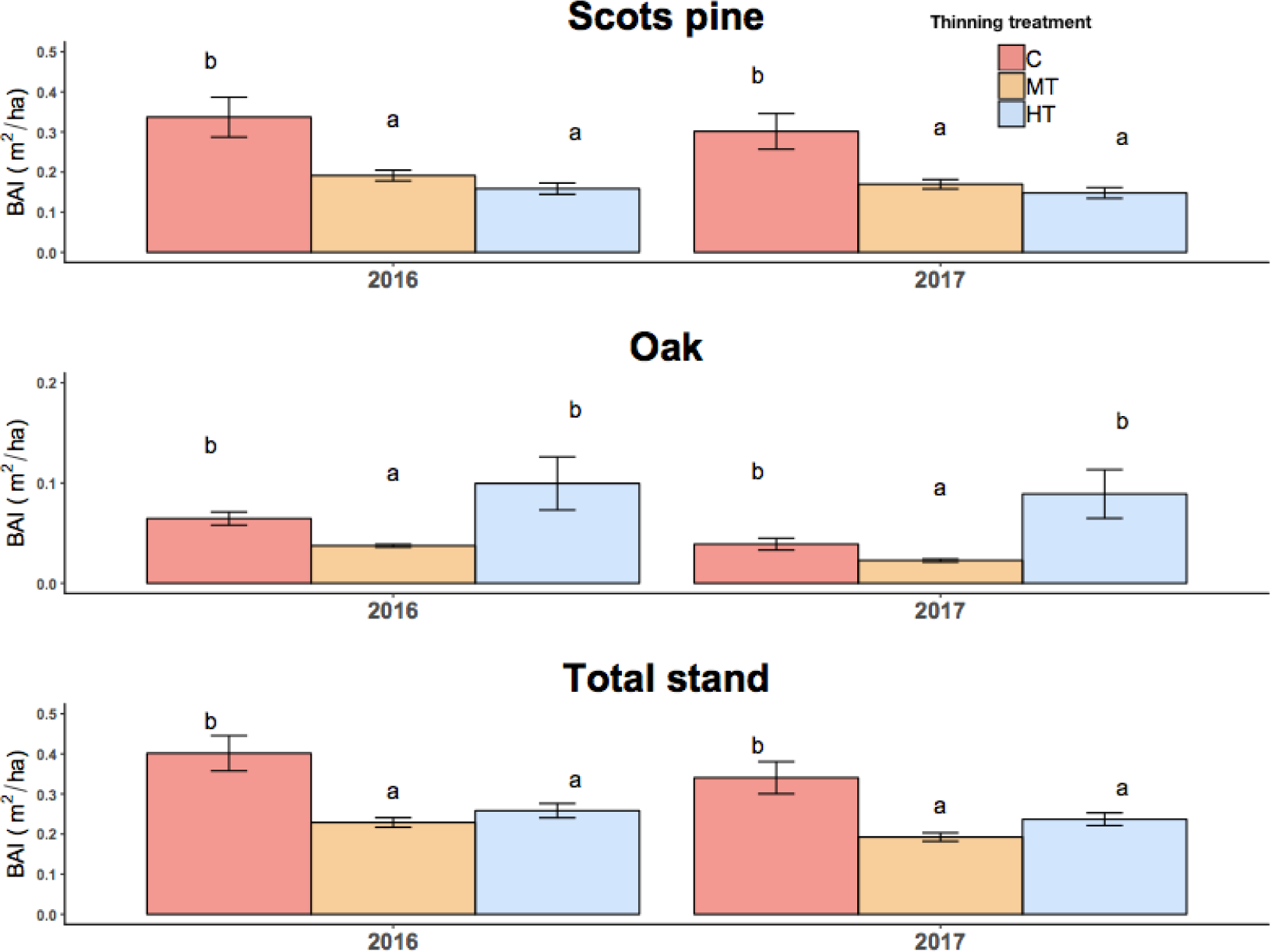
Unthinned and thinning treatments differences in basal area increment at stand level for each species and year. Total stand denotes the sum of basal area increment for both species. ‘C’ indicates unthinned, ‘MT’ moderate thinning and ‘HT’ heavy thinning treatments. Error bars show standard error and letters significant differences (P<0.05) between treatments for each year.

## 4. Discussion

### 4.1. The species growth differences

No difference was observed in the asymptote parameter between species during 2016 (which could be considered as mean spring weather conditions year), but only tree size (i.e., higher mean diameter for pine) and thinning treatment drove radial increment (Table 2). The absence of species differences under normal weather conditions could be consequence of both species are well adapted to the usual site conditions. However, the climate conditions in 2017, low precipitation in March and April (Figure 1), resulted in a reduction of spring asymptote for oak trees, but an increase for Scots pine (Table 2). Scots pine trees could take advantage of soil water reserves during spring droughts due leaf budburst and foliation differences: oak is still leafless in early spring and lives from the reserves of the previous year (Bauhus et al., 2017b; Fernández-De-Uña et al., 2017). Consequently, different timing in water use may cause that, when oak photosynthesis begins, scarce water is available due to it was largely consumed by Scots pine. As result, oak could be more sensitive to soil water deficits in spring, while Scots pine may have a higher sensitivity to summer droughts (Fernández-De-Uña et al., 2017; Merlin et al., 2015; Steckel et al., 2020). *Q.pyrenaica* is characterized by a complex clonal shallow root system with most coarse roots growing horizontally within a soil layer of one-meter depth, and rare vertical roots growing below (Salomón et al., 2016).

Drought tolerance could drive also species differences in inflection point (Table 2). A tighter stomatal control and higher intrinsic water use efficiency for Scots pine compared to oak may restrict growth earlier (Fernández-De-Uña et al., 2017). Early inflection point for Scots pine trees could involve a greater resource expenders and develop riskier life strategies to capture resources (Cuny et al., 2012), although it may be also an advantage to cope to drought stress by consuming firstly the scarce supplies of soil water (Fernández-De-Uña et al., 2017).

Species differences were also found for spring radial increment rate, with higher rates for Scots pine than oak (Table 2). Pioneer or early successional species, as Scots pine compared to oak, have light-demanding traits which allow to increase quickly radial growth (Sánchez-Costa et al., 2015). On the other hand, shade-tolerant oak trees would utilize resources more efficiently and could result even in higher radial increment rates (Cuny et al., 2012). Conversely, we observed lower increment rate for oak trees which could be caused by differences in reproduction methods (seed vs. coppice). Clonal structure influences stem growth in oak coppices, so bigger individual stool (clonal clump) is less vigorous (Salomón et al., 2013), which also difficulties the oak coppice management (Salomón et al., 2017). In addition, over-aging could affect also growth rates and causes a steep reduction in latewood production, acting as a predisposing factor in decline episodes of oak coppice stands (Corcuera et al., 2006). As a result, radial increment is mostly higher for pine than coppice oak trees in Mediterranean mixed forest (Aldea et al., 2018, 2017).

Finally, the influence of year factor on autumn asymptote (Table 2) suggest a probably diameter swelling/shrinkage in absence of growth. Camarero et al., (2010) and Fernández-De-Uña et al., (2017) reported that cambial reactivation, typical of Mediterranean species (Pacheco et al., 2018; Vieira et al., 2015), was commonly absent in both species studied here. Consequently, the contraction effect during 2017 autumn was similar for both species (Figure 2 and Table 2). This behavior is unusual since Mediterranean tree species generally experiment stem rehydration in autumn season after summer shrinkage caused by water deficit (Albuixech et al., 2012; Sánchez-Costa et al., 2015).

Overall, the species differences observed could be caused by species-specific physiological traits resulting in temporal and space growth diversification, which may relax resource competition by complementary exploitation of light and water (Williams et al., 2017). In this way, species mixing may involve a complementary and efficient use of resources over time by facilitation or competition reduction (Forrester, 2014). Thereby, del Río and Sterba, (2009) found that Scots pine-oak mixed stands support higher volume increment per occupied area compared to pure stands, similar as in central Europe for Scots pine mixed with sessile oak (Pretzsch et al., 2020; Steckel et al., 2019).

### 4.2. Thinning effect at different scales

Our results showed that thinning caused a clear trade-off between tree and stand growth. Heavy thinning affected both species at tree level increasing the spring asymptotic value (Table 2 and Figure 2), with no evidences to prolong the growing period (van der Maaten, 2013), although this result could change with higher temporal resolution in the measurements as recorded by electronic continuous dendrometers. Although some studies report a lack of response to thinning in larger trees (Varmola et al., 2004), thinning effect was independent of tree size, probably because it was even-aged stand or due to mechanisms related to resources partitioning for the tree species involved. We also found that heavy thinning had a positive effect at tree level regardless year or weather conditions (Table 2). Thinning has already been proved to reduce competition among trees, increase resources availability and, consequently, promote growth for the remaining trees for both species studied here (Cañellas et al., 2004; del Río et al., 2017a, 2008). However, the positive heavy thinning effect compared to moderate intensity seemed to be more evident during 2017 for both species (Figure 3). This may be due to the tree growth response to competition reduction could be delayed (Pukkala et al., 2002), caused by a great decompensation between subterranean and aerial parts of trees in the first year after thinning (Vincent et al., 2009). In addition, spring drought conditions during 2017 could affect growth response to thinning intensities. This result agrees with several studies which confirm that increasing the availability of growing space through thinning can enhance tree resistance and resilience in terms of height, diameter, and volume growth (D’Amato et al., 2013; Rais et al., 2014; Sohn et al., 2016b). Trees under low competition would better withstand the warmer conditions but with species-specific responses (Fernández-de-Uña et al., 2015). In this way, Sohn et al., (2016a, 2016b) found that heavy thinning improves growth recovery following drought events for Scots pine and, similarly, Cotillas et al., (2009) showed that selective thinning improved tree oak growth in stands under reduced rainfall conditions. Heavy thinning resulted in the greatest annual radial increment for pine even during a drought event in a Mediterranean pine-oak mixed forest (Aldea et al., 2017).

The absence of differences between unthinned and moderate thinning at tree level could be due to the initial spatial heterogeneity, i.e., local tree competition of target trees with band dendrometers could be similar for both treatments (unthinned and moderate thinning), and the short time elapsed after the thinning. A thinning effect analysis based on intra- and inter-specific tree competition index and longer period may clarify the results obtained here and could be the aim for a future work. However, although it was not revealed by the fitted model, moderate thinning could increase also tree radial increment for oak compared to unthinned stands (Figure 3). This agrees with Corcuera et al., (2006), who show how thinned oak trees formed wider tree-rings, more latewood and multiseriate tree-rings than overaged trees. On the other hand, historical oak coppycing management could cause high carbon expenditures in root respiration and centennial clones may constrain their aboveground development, so thinning might not succeed or even enhance a physiological root-to-shoot imbalance (Salomón et al., 2017).

Our results showed that total stand basal area increment declines with thinning, indicating a loss of productivity in thinned stands in comparison to unthinned stands. However, there was not significant difference between the two thinning intensities, in contrast to the typical decreasing pattern with strong density reductions (del Río et al., 2017b; Pretzsch, 2020). When focusing on species level, the lower Scots pine basal area increment in thinned stands agrees with the findings for this species in monospecific stands (del Río et al., 2008), although the magnitude of growth loss depends on the region, site and stand age (del Río et al., 2017a). For oak, our results showed that basal area increment can be maintained or even increased with heavy thinning, but moderate thinning reduced the growth. Cañellas et al., (2004) reported a higher tree growth under heavy thinning treatment for coppice oak monospecific stands, although no differences were found for stand yield (total basal area and biomass). The species specific responses to thinning explain the pattern at stand level, as the greater oak basal area increment in heavy thinning reduces the differences between the control and heavy thinning treatments (Fig. 4). This suggests that species mixtures may reduce the loss of productivity due to thinning in comparison with monospecific stands and that their growth response pattern differs from those of monospecific stands (Thurm and Pretzsch, 2021).

Accordingly, Pretzsch, (2003) reported lower increment losses for mixed stands of Norway spruce-beech than in pure stands when density was reduced. Primicia et al., (2016) even observed that stand volume annual increments were higher in the thinned than in the unthinned plots in a Scots pine-beech mixed stand where only pine trees were removed. However, heavy thinning in dry sites, such as the studied here, may generate negative effect on stand growth by canopy opening which outweighs the more efficient use of soil resources as water availability. Anyway, our results at short term evidenced that stand production decreases with thinning intensity (Ashton and Kelty, 2018), causing stand growth loss and consequently a reduction in carbon sequestration. Therefore, thinning caused a marked trade-off between tree and stand growth levels, although the positive outcomes could offset the negative ones, depending on the target of forest management (Vilà-Cabrera et al., 2018). The greatest positive and most obvious effect of heavy thinning is to enhance oak growth independently of scale. These structural changes could lead to a new management strategy towards a conversion from overaged coppice stands into high forests (originating from seed), conforming a stand growth stability from the mixture (del Río et al., 2017b). Other ecosystem functions and services may be favored as a fire risk reduction, biodiversity conservation or an improvement of conditions for oak fruiting and sexual regeneration. In addition, remain trees of Scots pine would increase the size for final harvest and, hence, the wood quality and timber value.

## 5. Conclusions

The implementation of a given forest management practice implies the existence of trade-offs among forest functions responses, i.e., it may be beneficial for reaching a specific objective but, at the same time, it can impair the consecution of other objectives or induce negative impacts on untargeted ecosystem components (Vilà-Cabrera et al., 2018). Here, we showed that, although thinning supposes a marked trade-off between tree and stand production in the mixture, oak trees can overcome it and be favored. Our results evidence a good strategy towards a conversion from overaged coppice stands into high forests, conforming a stable mixed forest stand. Besides, higher growth for Scots pine residual trees could also cause an increase in timber value or be beneficiated by a competition release of water resources, increasing simultaneously the resistance to droughts. Therefore, reducing the stand growth via heavy thinning to favor the dominated oak coppice trees may be a reasonable management proposal to ensure the stability and persistence of this type of mixture, which could play an important role for the ecosystem services preservation into a climate change scenario.

## Acknowledgments

This study has received funding from the European Union’s Horizon 2020 research and innovation programme under grant agreement No952314 and under the Marie Skłodowska-Curie grant agreement No 778322. Thanks for the support by the Spanish Ministerio de Ciencia e Innovación (# PID2021-126275OB-C21/C22). Thanks go furthermore to the Junta de Castilla y Leon, Spain, and the European Union for funding the Projects VA183P20 (SMART— Bosques mixtos: Selvicultura, Mitigacion, Adaptacion, Resiliencia y Trade-offs) and CLU-2019-01—iuFOR Institute Unit of Excellence of the University of Valladolid through the ERDF “Europe drives our growth”. Jorge Aldea’s work was funded by a partnership grant of the University of Valladolid and Banco Santander, and the grant RYC2021-033031-I, funded by MCIN/AEI/10.13039/501100011033 and by the European Union “NextGenerationEU”/PRTR”. The authors acknowledge all those who participated in the plot design installation and implementation of tree measurements, especially to the Forestry Services of the Leon province, where the sampling took place. The authors thank the Spanish State Meteorological Agency (AEMET) of the Ministry of Agriculture, Food and Environment for granting access to the meteorological data.

## Appendix A. Supplementary Figures

**Figure S1.**
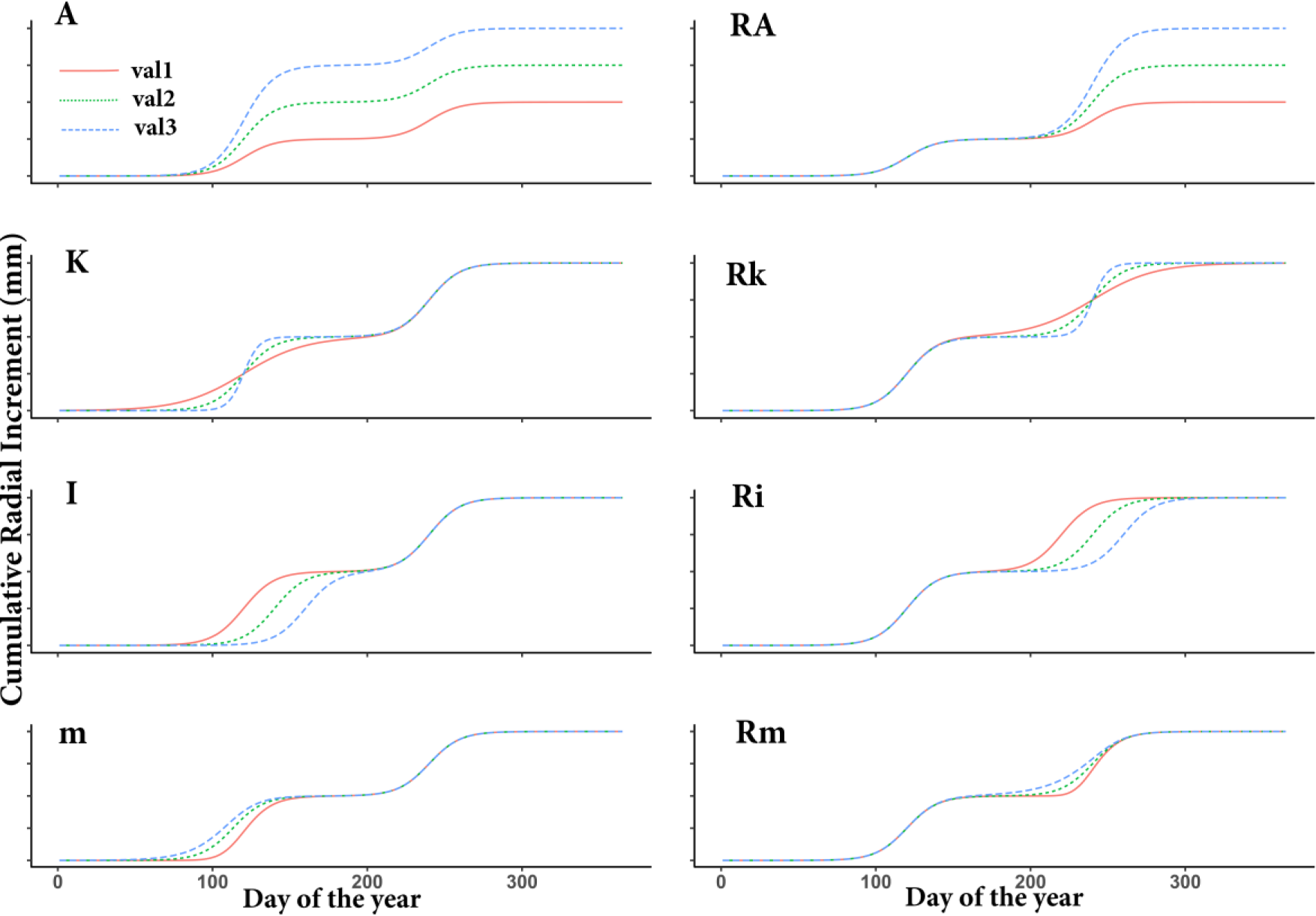
Shift value display in parameter (val1<val2<val3) for the double-Richards model (Eq.1).

