## Supplementary material for "Short-term thinning effect on inter- and intra-annual radial increment of Mediterranean Scots pine-oak mixed forest": Figure S1

### Supplementary Figures

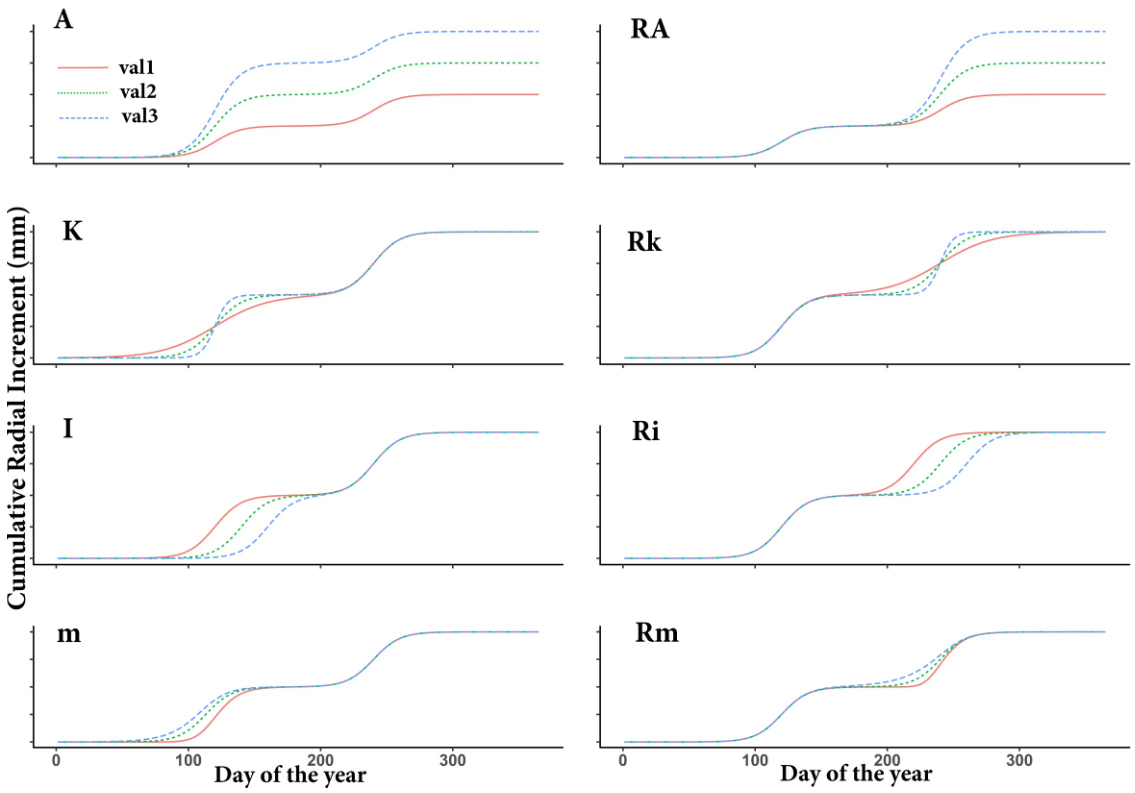


*Figure S1. Shift value display in parameter (val1<val2<val3) for the double-Richards model (Eq.1).*
